# Meta-analysis of the human brain transcriptome identifies heterogeneity across human AD coexpression modules robust to sample collection and methodological approach

**DOI:** 10.1101/510420

**Authors:** Benjamin A. Logsdon, Thanneer M. Perumal, Vivek Swarup, Minghui Wang, Cory Funk, Chris Gaiteri, Mariet Allen, Xue Wang, Eric Dammer, Gyan Srivastava, Sumit Mukherjee, Solveig K. Sieberts, Larsson Omberg, Kristen D. Dang, James A. Eddy, Phil Snyder, Yooree Chae, Sandeep Amberkar, Wenbin Wei, Winston Hide, Christoph Preuss, Ayla Ergun, Phillip J Ebert, David C. Airey, Gregory W. Carter, Sara Mostafavi, Lei Yu, Hans-Ulrich Klein, the AMP-AD Consortium, David A. Collier, Todd Golde, Allan Levey, David A. Bennett, Karol Estrada, Michael Decker, Zhandong Liu, Joshua M. Shulman, Bin Zhang, Eric Schadt, Phillip L. De Jager, Nathan D. Price, Nilüfer Ertekin-Taner, Lara M. Mangravite

## Abstract

Alzheimer’s disease (AD) is a complex and heterogenous brain disease that affects multiple inter-related biological processes. This complexity contributes, in part, to existing difficulties in the identification of successful disease-modifying therapeutic strategies. To address this, systems approaches are being used to characterize AD-related disruption in molecular state. To evaluate the consistency across these molecular models, a consensus atlas of the human brain transcriptome was developed through coexpression meta-analysis across the AMP-AD consortium. Consensus analysis was performed across five coexpression methods used to analyze RNA-seq data collected from 2114 samples across 7 brain regions and 3 research studies. From this analysis, five consensus clusters were identified that described the major sources of AD-related alterations in transcriptional state that were consistent across studies, methods, and samples. AD genetic associations, previously studied AD-related biological processes, and AD targets under active investigation were enriched in only three of these five clusters. The remaining two clusters demonstrated strong heterogeneity between males and females in AD-related expression that was consistently observed across studies. AD transcriptional modules identified by systems analysis of individual AMP-AD teams were all represented in one of these five consensus clusters except ROS/MAP-identified Module 109, which was specific for genes that showed the strongest association with changes in AD-related gene expression across consensus clusters. The other two AMP-AD transcriptional analyses reported modules that were enriched in one of the two sex-specific Consensus Clusters. The fifth cluster has not been previously identified and was enriched for genes related to proteostasis. This study provides an atlas to map across biological inquiries of AD with the goal of supporting an expansion in AD target discovery efforts.

## INTRODUCTION

Alzheimer’s Disease (AD) is a debilitating neurodegenerative disease affecting more than 5 million Americans for which we lack effective long-term disease-modifying therapeutic strategies (Cummings et al., 2014). Several therapeutic mechanisms are under active evaluation in clinical trials (Kumar et al., 2015) across the field - including the amyloid hypothesis. Because AD is likely to result from molecular dysregulation across a series of biological systems within the brain (De Strooper and Karran, 2016), there is some question as to whether therapeutic targeting of a single pathway will be sufficient to completely address the full burden of this disease. Furthermore, recent evidence suggests that AD may be a collection of conditions with multiple underlying causes that lead to similar symptomatic and pathological end points (Brenowitz et al., 2017; Winblad et al., 2016). For these reasons, there is need to pursue a diverse set of mechanistic hypotheses for therapeutic intervention.

Systems biology analysis can provide a rich mapping of the inter-related molecular dysregulations involved in AD that may be useful to guide drug target discovery towards a diverse set of complementary therapeutic mechanisms. Several systems-level analyses of human AD brain have been previously reported (Allen et al., 2018a, 2018b; Lu et al., 2014; Mostafavi et al., 2018; Seyfried et al., 2017; Zhang et al., 2013a). The importance of neuroinflammation in AD has been described across these (Patrick et al., 2017; Zhang et al., 2013a) and also from genetic studies (Carrasquillo et al., 2017; Efthymiou and Goate, 2017; Guerreiro et al., 2013; Jin et al., 2015; Jonsson et al., 2013; Raj et al., 2014; Sims et al., 2017), supporting a major focus on this pathway for therapeutic development as is currently underway (Ardura-Fabregat et al., 2017). In addition to neuroinflammation, other molecular pathways have been identified from systems biology studies of both RNA and protein abundance including multiple processes related to oligodendrocytic functions such as myelination (Allen et al., 2018a; McKenzie et al., 2017; Mostafavi et al., 2018; Seyfried et al., 2017). In several cases, these compelling observations arise from individual studies and, as of yet, their reproducibility is unknown.

This project aims to develop an atlas of AD-associated changes in molecular state that provides a mechanism to evaluate the consistency and robustness of systems analyses and the use of their findings to support AD target discovery. To this aim, we build on the resources and expertise gathered across the Accelerating Medicines Partnership in Alzheimer’s Disease Target Discovery and Preclinical Validation project (AMP-AD – ampadportal.org). AMP-AD focuses on identification of AD disease drivers using systems-level evaluation of disease state in human brain tissue. A major early outcome of this consortium was the generation and public release of RNA-seq data generated across three sizable but distinct human postmortem brain studies that are distributed through the AMP-AD Knowledge Portal – https://ampadportal.org (Allen et al., 2016; Jager et al., 2018; Wang et al., 2018). Here, we use coexpression meta-analysis across these studies to develop a robust systems-level molecular atlas of AD. Coexpression network analysis is a commonly used data-driven approach to identify gene sets (or modules) that are similarly co-expressed across samples in a data set (Langfelder and Horvath, 2008a). These modules are often comprised of genes involved in biological processes that interact and/or exhibit coordinated activity in response to molecular and cellular states, pathological processes, and other factors (Gaiteri et al., 2015). Although distinct and important biology may be uniquely represented in any one of these studies, meta-analysis provides a generalized illustration of the changes in transcriptional state associated with AD in a manner that is robust to technical confound and study heterogeneity. An atlas derived of cross-study AD-associated transcriptional modules can support target discovery by (1) promoting target discovery across multiple distinct biological processes, (2) informing experimental design for target validation studies, (3) creation of improved experimental models and assessment of current experimental models (Wan et al., submitted), and (4) evaluating population heterogeneity in disease pathophysiology that may impact therapeutic efficacy.

## RESULTS

### AMP-AD collection of human RNA-seq data

We analyzed existing transcriptional data generated from post-mortem brain tissue homogenate from three separate sample sets including the Religious Order Study and the Memory and Aging Project (ROSMAP) (Bennett et al., 2012b, 2012a; Jager et al., 2018), the Mount Sinai Brain Bank (MSBB) RNA-seq study (Wang et al., 2018), and the Mayo RNA-seq study (Allen et al., 2016) (Mayo). Samples were collected from seven distinct brain tissues - dorsolateral prefrontal cortex (DLPFC) in ROSMAP; temporal cortex (TCX) and cerebellum (CBE) in Mayo, and inferior frontal gyrus (IFG), superior temporal gyrus (STG), frontal pole (FP), and parahippocampal gyrus (PHG) in MSBB. Several differences in data collection and processing protocols across studies were identified and accounted for during data processing and analysis (see **Table 1** and **Methods**).

### Development of AD-related transcriptional modules by consensus coexpression network analysis

To identify AD-related human transcriptional modules that were robustly observed in a generalized manner across methods and studies, we performed a consensus analysis for all seven tissue types using five coexpression analysis methodologies. These five distinct coexpression learning algorithms included: MEGENa (Song and Zhang, 2015), rWGCNA (Parikshak et al., 2016), metanetwork (**Methods**), WINA (Wang et al., 2016) and SpeakEasy (Gaiteri et al., 2015). Independent performance of each of the 5 methods across each of the 7 tissues identified 2,978 tissue-specific coexpression modules (CBE: 458, DLPFC: 450, FP: 393, IFG: 429, PHG: 370, STG: 336, TCX: 502, 10.7303/syn10309369.1). As expected, similar coexpression structure was observed in each data set across methods, as indicated by significant overlap in module memberships (**Figure S1**). Within each tissue, we next identified AD-related modules that were well-represented across analysis methodologies. This analysis was limited to those modules that were significantly enriched for differentially expressed genes related to AD based on a meta-analysis of differential expression across the seven brain regions (**Methods**). To do this, graph clustering (Pons and Latapy, 2005) was performed on all modules within a tissue with an edge betweenness community identification algorithm (Pons and Latapy, 2005) and weights from the Fisher’s exact test estimate (**Methods, Figure S1**). This meta-analysis of coexpression modules and differential expression signatures identified 30 AD associated modules across the seven tissue types (CBE: 4, DLPFC: 4, FP: 4, IFG: 4, PHG: 5, STG: 4, TCX: 5, 10.7303/syn11932957.1).

To establish confidence that these consensus modules provided an improved and coherent representation of AD-altered biology, they were evaluated for enrichment of gene sets previously identified as relevant to AD (**Figure 1** and **Methods**). The consensus modules showed an improved percentage enrichment for previously published AD related gene sets or pathways relative to (a) the differential expression meta-analysis gene set (P-value = 0.036, Wilcoxon rank sum test), (b) the modules defined by individual coexpression methods (P-value < 2x10^-16^ Wilcoxon rank sum test) and (c) those modules defined previously in the literature (Zhang et al., 2013b) (P-value 3.9x10^-13^ Wilcoxon rank sum test) (**Figure 1**). Evaluation across tissues demonstrated that these 30 consensus modules fell into five well-defined clusters that were highly preserved across study and tissue type and demonstrated a low degree of overlap between clusters (**Figure 2A**) based on a Fisher’s exact test of gene membership overlap. These five ‘consensus clusters’ were used to represent distinct patterns of AD-related transcriptional state that were consistently observed.

**Figure 1.**
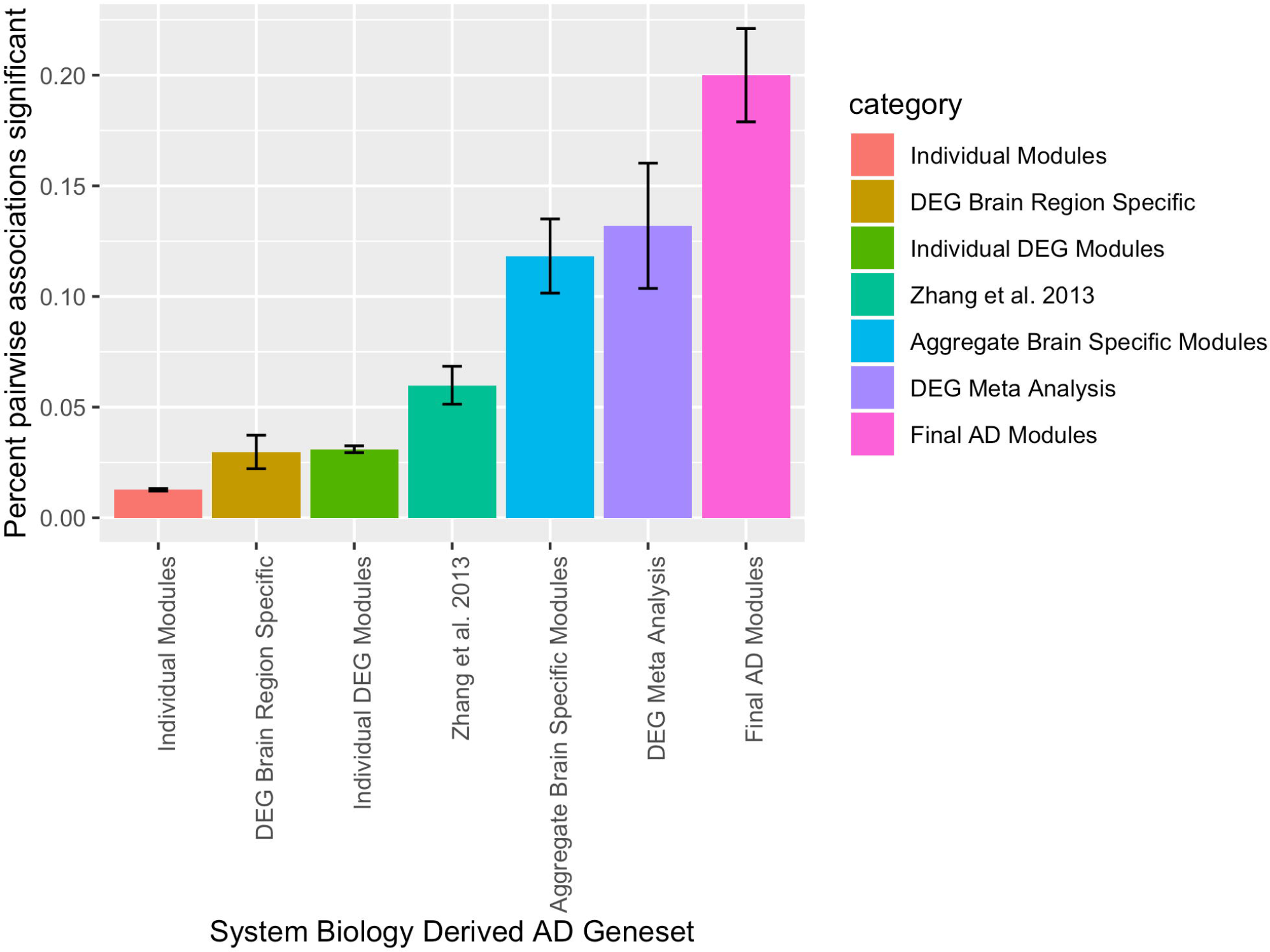
Percentage of total pairwise module by literature curated AD gene set associations significantly enriched (FDR <=0.05) for 12 known AD gene sets with standard errors shown.

**Figure 2.**
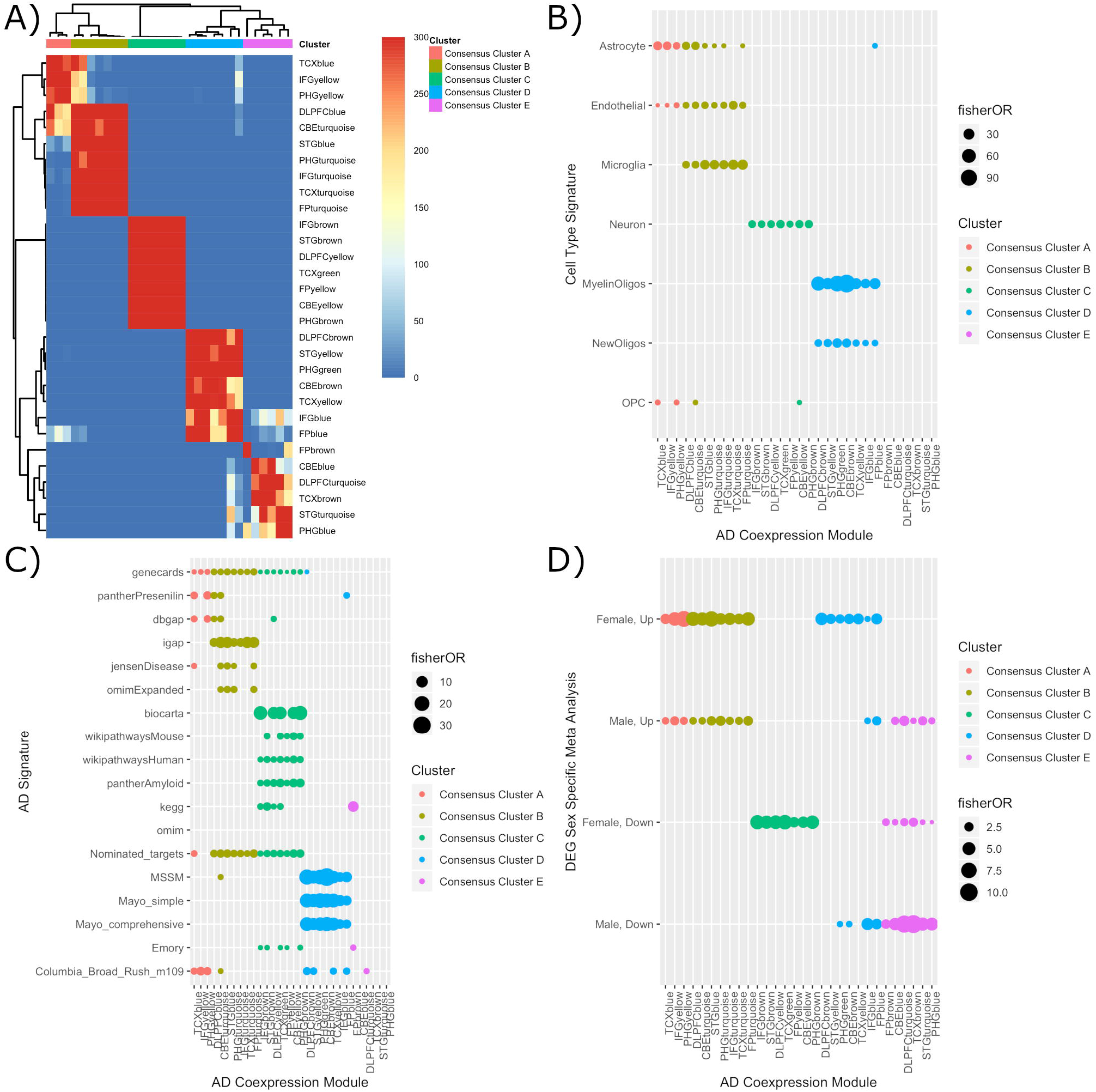
A) Overlap between identified 30 AD associated coexpression modules - there are 5 predominant clusters identified across brain regions – Consensus Cluster A, B, C, D, and E. B) Cell type enrichments of the 30 AD associated coexpression modules. C) Enrichment for curated AD gene sets within the 30 AD associated coexpression modules. D) Enrichments for differentially expressed genes based on the meta-analysis in the 30 AD associated coexpression modules.

Because previous studies have identified cell-type specific changes from transcriptional analysis, we next evaluated enrichment of cell-type specific gene signatures as identified from previously published cell type specific RNA-seq data from Zhang et al (Zhang et al., 2014). While most of the genes represented in these consensus clusters were not cell type specific, we did observe that genes represented in cell-type specific gene signatures were grouped by cluster. As shown in **Figure 2B**, cell-type specific gene sets clearly clustered into four of five consensus clusters: the astrocytic signature was enriched in Consensus Clusters A and B, the endothelia and microglial signatures were enriched in Consensus Cluster B, the neuronal signature was enriched in Consensus Cluster C, and the oligodendroglial signature was enriched in Consensus Cluster D. While we see significant enrichment for cell type specific gene-sets in Consensus Cluster A-D, these modules are large (2090 ± 1150 genes, **Table S3**), and show enrichment for a diverse set of biological processes beyond cell type specific processes (10.7303/syn11954640.1). Accordingly, Consensus Cluster E was not enriched for cell type specific signatures, but instead was consistently enriched for genes that were associated with proteostasis – including with the attenuation phase of the transcriptional response to heat shock (Abravaya et al., 1991; Fabregat et al., 2016), detection of unfolded protein (GO:0002235) (2015), response to unfolded protein (GO:0006986) (2015), and HSF1 activation (Cotto et al., 1996; Fabregat et al., 2016; Zuo et al., 1995) (**Table 2**).

### Heterogeneity in expression of consensus clusters between females and males

To evaluate whether consensus clusters may prove useful in the identification of molecular heterogeneity in disease across populations, we evaluated sex-specific differences across clusters in AD-related gene expression. Indeed, sex-specific differential expression gene (DEG) sets – from a sex specific meta-analysis of differential expression across the seven tissue types - were differentially enriched across the consensus clusters for females vs. males (**Figure 2D**). Consensus Clusters A and B demonstrated similar direction of DEGs across sex through enrichment was stronger in females vs. males. This suggests that transcriptional changes related to neuroinflammatory processes were common across the sexes, albeit more pronounced in females. In contrast, the Consensus Clusters C and D were strongly enriched for genes altered in females but not in males, suggesting that the overall association of these clusters with AD-related differential expression was predominantly driven by females. Consensus Cluster E was enriched for genes that were down-regulated in both male and female AD cases. This last cluster was moderately enriched for genes that were up-regulated in male AD cases, with no such enrichment observed in females. These changes demonstrate that sex-specific changes in AD-related gene expression are heterogenous across consensus cluster.

### Use of consensus modules as an atlas to evaluate diversity of AD target discovery efforts

The consensus clusters can be used in aggregate as an atlas to support the selection of a robust and diverse set of AD therapeutic hypotheses for target discovery. To this aim, we next evaluated the enrichment across clusters of AD biology under active investigation for target discovery (**Figure 2C**). Consensus Clusters A, B, and C were enriched for AD pathways derived from the scientific literature (Amberger et al., 2015; Kanehisa et al., 2017; Kutmon et al., 2016; Lambert et al., 2013; Mi et al., 2017; Nishimura, 2001; Safran et al., 2010; Tryka et al., 2014), AD gene sets (Lambert et al., 2013; Tryka et al., 2014) derived from genetic association analyses, and pathways related to therapeutic hypotheses currently undergoing active drug development including the amyloid secretase pathway (neuronal cluster), the Presenilin amyloid processing pathway (astrocytic cluster), and deregulation of *CDK5* pathway that is implicated in Tau hyperphosphorylation (neuronal cluster).

We next evaluated the consensus clusters for enrichment of modules identified for AD target discovery by the individual teams within the AMP-AD consortium, as part of their systems biology-based target discovery programs. Each team had identified AD-related modules through study-specific analyses (Allen et al., 2018a; Johnson et al., 2018; McKenzie et al., 2017; Mostafavi et al., 2018). We evaluated how the primary findings from these individual analyses mapped onto the consensus clusters. First, we examined enrichment the results published from independent analysis of the ROS/MAP study (Mostafavi et al., 2018). This original analysis identified several modules that were associated with rate of cognitive decline including module 109. Unlike the modules previously identified within the literature or by the other teams (see below), module 109 membership did not group into a single consensus cluster. Instead, module 109 membership was enriched across 4 of the 5 consensus clusters. Strikingly, module 109 was the most strongly enriched for genes that are up in AD cases (FET OR, P-value: 9.8, 2x10^-72^), up in male AD cases (FET OR, P-value 7.4, 1.1x10^-22^), and up in female AD Cases (FET OR, P-value: 9.2,2x10^-78^ **Table S5, Table S7, Table S8**) among all AD signatures we tested.

In contrast, the modules identified by the other AMP-AD teams were all enriched within a single consensus cluster. The Mayo and Mt Sinai teams each identified a separate module that was enriched for oligodendrocyte signatures and that significantly overlapped with consensus cluster D (**Figure 2C, Table S4**). Notably, AD-related decreases in expression of genes within these two modules were reported by each team. Because we observed sex-specific differences in AD gene expression in consensus cluster D but results from sex-specific analyses were not reported by either team, we evaluated the Mayo and Mt. Sinai modules for sex-specificity. Indeed, the sex-specific pattern of AD-related expression were also consistently observed in both the Mayo and Mt. Sinai modules. For the Mayo module, the effect size for AD-related increased expression in females was much smaller than the effect size for AD-related decreased expression in males (**Table S8, S10**), supporting a modest decrease in expression based on combined analysis across all AD cases as was reported by the Mayo team (Allen et al., 2018a). The Mt. Sinai module demonstrated the same AD-related increased expression in females but no significant changes in males. The genes represented in the Mt Sinai module may be specific to the female signature because Mt. Sinai module membership was derived from analysis of a sample set with a preponderance of female samples. Finally, we evaluated cluster enrichment for an RNA-binding module identified by the Emory AMP-AD team (Johnson et al., 2018) from systems analysis of proteomic data. This was enriched in the Consensus Cluster B suggesting that it is co-expressed with genes involved with synaptic function.

Finally, we evaluated enrichment for the one hundred genes nominated by the AMP-AD consortia as the first set of candidates for AD target evaluation (https://agora.ampadportal.org). Because these targets were selected in part based on analysis of these data, we expected to observe significant enrichments within the consensus clusters. Interestingly, significant enrichment was observed (adjusted P-value < 0.05), but this was specific to Consensus Clusters A, B, and C – those that were also enriched for previously known AD biology processes. This suggests that the initial round of AMP-AD target nominations was guided by data-driven analysis in combination with evaluation of prior biological knowledge. Subsequent nominations would benefit from an expansion into biology represented within the other consensus clusters that are equally robust but have been less studied in the context of AD – particularly biology within Consensus Cluster D that was observed across multiple independent analyses.

## DISCUSSION

Identification of therapies for the treatment or prevention of AD has been hampered by many difficulties including a limited pipeline of well-validated targets (Kumar et al., 2015). In part, this is because target discovery and validation has been plagued by multiple issues including: (a) poor understanding of the complex inter-related biological processes that are dysregulated with AD on a systems level (De Strooper and Karran, 2016), (b) presence of other aging-related neuropathologies that confound interpretation of differential expression studies(De Jager et al., 2018), (c) evaluation of AD biology in experimental model systems that do not effectively recapitulate human disease (King, 2018), and (d) reliance on therapeutic hypotheses that show no efficacy in late stage clinical trials (Makin, 2018). Using molecular data collected from human brains, this project provides an overview of the systems-level models of AD state in human brain that can be used to inform identification and assessment of complementary target hypotheses. Human brain transcriptional data collected from three cohorts were used to define a robust, reproducible set of human AD associated coexpression modules by consensus network analysis across five co-expression network methodologies. Consensus modeling provided a set of generalizable observations that robustly define AD-associated dysregulation in transcriptional state, and as such provide a resource to guide target selection and validation strategies.

The consensus analysis identified 5 consensus clusters that represented distinct patterns of AD-related changes in gene expression. The observed AD-associated changes in transcriptional state were consistently observed across all brain regions except cerebellum in terms of the differential expression patterns (the coexpression patterns are conserved). The reason why these signatures are not observed with AD in cerebellum is unknown but may be caused by region-specific differences in AD-associated transcriptional dysregulation, in AD pathology, and/or in cellular resilience to AD pathology (Stowell et al., 2018). The interpretation of these differences is also confounded by the basic differences between cerebellar cortex and cerebral cortex in terms of cell composition, cellular architecture, and function (von Bartheld et al., 2016).

These consensus clusters provide a general framework for evaluating heterogeneity in disease across populations. In this analysis, few observed a drastic difference in AD-related expression changes in females vs. males within each consensus cluster. For 4 of 5 Consensus Clusters, females exhibited significantly greater AD-associated expression changes as compared to males. This included greater increases in expression of the module clusters that were enriched for astrocytic, microglial, endothelial, and oligodendroglial signatures and greater reduction in expression of those enriched for neuronal signatures. In contrast, AD-associated alterations in expression of Consensus Cluster E was more prominent in males than in females. This cluster was enriched for response to unfolded protein and heat shock response.

Increasingly, the literature supports differences between males and females in AD progression, although it is unknown whether these are caused by differences in AD-mediated processes, in rate of progression within comparable processes, in pre-disease state or in some other cause (Mielke et al., 2014; [CSL STYLE ERROR: reference with no printed form.]). Furthermore, we see evidence of sex specific differences in genetic regulation of disease (Nazarian et al., 2018), including at the level of expression of the oligodendrocyte myelinating cell module. A suggestive hypothesis is that AD genetic loci identified to date are highly enriched in neuroinflammatory modules *precisely* because they show similar biology (at least at the transcriptomic level) in both men and women (**Figure 2D**). This indicates that genetic association analyses stratified by sex may further illuminate some of the missing genetic factors underlying Alzheimer’s disease. Preliminary evidence suggests that this may be the case: a genetic risk score calculated in ROSMAP based on the 21 IGAP risk loci was associated with an eigengene in the oligodendrocyte consensus cluster (specifically the DLPFC brown module) (adjusted p-value, Bonferroni: 10^-2^), but only in females and was significantly different from males. Further disentangling the role of sex and genetics in understanding disease heterogeneity will be key to development of efficacious therapeutic interventions, especially if there are different underlying mechanisms driving disease etiology between men and women.

Identification of conserved human AD-related consensus clusters provides several benefits in the pursuit of AD target discovery. First, they highlight the distinct aspects of brain biology that are dysregulated with AD– including several that are not currently under active investigation for drug development. While not all dysregulated pathways are likely to be causative, these results suggest that a broader range of therapeutic hypotheses exist and serves to guide researchers to areas of biology that may merit further pursuit. In this manner, the consensus clusters were used to demonstrate three major areas of AD-related biology that are under active pursuit for drug discovery and a fourth area of interest. This fourth, represented by Consensus Cluster D, had a complex subcluster architecture that may contain several biological processes of interest for further pursuit. Indeed, two of these subclusters were independently identified and are under active pursuit by the AMP-AD Mayo and Mt. Sinai target discovery teams respectively. We note that the use consensus methodologies is explicitly designed to identify signatures of disease that are most robust to technical and study specific heterogeneity and, as such, can provide evidence to support costly drug discovery programs. This does not preclude the relevance of other interesting biology that was not recapitulated across studies due to small effect size or uneven representation across studies based on differences in sample ascertainment.

In addition to evaluating diversity in AD therapeutic hypotheses, these human AD-related clusters can be used to identify appropriate experimental model systems for further evaluation. Traditionally, AD model systems have been developed through genetic perturbations of one or more AD-related pathways (King, 2018). While none of these models provides a complete recapitulation of human disease, many provide a useful framework to evaluate dysregulation within specific pathways. Since a subset of human co-expression clusters display conserved co-expression and/or overlapping differential expression in brains from AD mouse models, these human clusters may help to assess the appropriateness of AD experimental models for pathway-specific evaluation – as well as to highlight other genetic perturbations that may provide useful model systems to complement those that are commonly used(Mostafavi et al., 2018; Neuner et al., 2018, Wan et al. submitted).

This analysis provides an important first step in developing a molecular framework to evaluate and promote diversity in AD target discovery. There are a handful of caveats to the approach taken in this study. First and foremost, this study focuses on transcriptional measures of disease response in post-mortem brain and, as such, provides an initial but incomplete picture of the molecular response and triggers of disease – including proteomic, epigenomic, and metabolomic signatures of disease. Previous work indicates the correlation between transcriptomic and proteomic signatures of disease is relatively modest (Pearson’s r = 0.30) (Seyfried et al., 2017), and thus a more thorough integrative analysis is warranted to determine the full space of molecular signature of disease progression. Additionally, all of the samples were from post-mortem tissue which could potentially introduce non-AD specific effects due to the state of the person at death (e.g. the effect of agonal state or preterminal decline in cognition immediately prior to death). Because we adopted a case/control analytic strategy to enable a meta-analysis across the three sources of data, each of which has a very different study design, we could not consider individuals with intermediate phenotypes. As such, this analysis is limited to a syndromic diagnosis of pathologic AD, further refined by including cognitive evaluations for the ROSMAP and MSSM subjects. Given limitations of available neuropathologic phenotypes, we were not able to consider the possible impact of other aging-related pathologies on our results. Finally, because these studies focused on whole tissue analysis, we cannot resolve which observed changes are driven by differences in the cellular composition of the tissue samples between AD cases and controls (neuronal death and reactive gliosis), and which are due to actual differences in the cellular expression levels. In looking for consensus modules across multiple brain regions that are variably influenced by AD pathology and further characterizing these modules based on additional evidence for involvement in AD, we have identified robust changes that may not entirely be driven by the former. However further work is needed to refine the molecular changes and pathways associated with AD and the implications for specific central nervous system cell-types.

These transcriptional AD-related module clusters represent an attractive mechanism to support translational research. Predictions of genes with an important role within an AD-related module cluster have been validated experimentally *in vitro* and *ex vivo* (Mostafavi et al., 2018; Yu et al., 2018; Zhang et al., 2013b). Within model systems, gene signatures for human AD clusters can serve as readouts to evaluate consequences of target engagement that are known to be relevant to human disease (Wan et al., submitted). Such experiments could also identify biochemical signatures – or consequences –associated with changes in human AD clusters that could be used to advance therapeutic hypotheses or identify endophenotypic biomarkers. While effectiveness of such approaches needs to be tested, such approaches are already underway in several programs including those using mouse, fly, and cell-based model systems to evaluate AD biology. An integrated, systems approach to AD target evaluation is a powerful opportunity to advance the field.

## ACKNOWLEDGEMENTS

The results published here are in whole or in part based on data obtained from the AMP-AD Knowledge Portal (<u>doi:10.7303/syn2580853</u>). ROSMAP Study data were provided by the Rush Alzheimer’s Disease Center, Rush University Medical Center, Chicago. Data collection was supported through funding by NIA grants P30AG10161, R01AG15819, R01AG17917, R01AG30146, R01AG36836, U01AG32984, U01AG46152, the Illinois Department of Public Health, and the Translational Genomics Research Institute. Mayo RNAseq Study data were provided by the following sources: The Mayo Clinic Alzheimer’s Disease Genetic Studies, led by Dr. Nilufer Ertekin-Taner and Dr. Steven G. Younkin, Mayo Clinic, Jacksonville, FL using samples from the Mayo Clinic Study of Aging, the Mayo Clinic Alzheimer’s Disease Research Center, and the Mayo Clinic Brain Bank. Data collection was supported through funding by NIA grants P50 AG016574, R01 AG032990, U01 AG046139, R01 AG018023, U01 AG006576, U01 AG006786, R01 AG025711, R01 AG017216, R01 AG003949, NINDS grant R01 NS080820, CurePSP Foundation, and support from Mayo Foundation. Study data includes samples collected through the Sun Health Research Institute Brain and Body Donation Program of Sun City, Arizona. The Brain and Body Donation Program is supported by the National Institute of Neurological Disorders and Stroke (U24 NS072026 National Brain and Tissue Resource for Parkinson’s Disease and Related Disorders), the National Institute on Aging (P30 AG19610 Arizona Alzheimer’s Disease Core Center), the Arizona Department of Health Services (contract 211002, Arizona Alzheimer’s Research Center), the Arizona Biomedical Research Commission (contracts 4001, 0011, 05-901 and 1001 to the Arizona Parkinson’s Disease Consortium) and the Michael J. Fox Foundation for Parkinson’s Research. MSBB data were generated from postmortem brain tissue collected through the Mount Sinai VA Medical Center Brain Bank and were provided by Dr. Eric Schadt from Mount Sinai School of Medicine. Furthermore, Emory study data were supported through funding by NIA grants P50 AG025688, U01 AG046161, and U01 AG061357.

## AUTHOR CONTRIBUTIONS

BAL, TMP, VS, MW, CF, CG, MA, PS, YC, CF, XW performed differential and network expression analyses. BAL, JE, KDD, PJE, PS performed bioinformatic analyses. BAL, TMP, CG, MW, LMM, SKS, KDD, PE, LMM designed the analysis plan. BAL, TMP, MA, NET, LMM, AL, DAB, PLDJ, JMS, GWC wrote the manuscript. BAL, TMP, LMM, KD, CG, MA, ED, GS, SM, SA, WH, HUK, CP, MD, KE, LY, AE, CP, GWC contributed to interpretation of analyses. DAC, TG, AL, DAB, KE, MD, ZL, BZ, ES, PLDJ, NDP, NET conceived the human study design.

## TABLE LEGENDS

**Table 1 -** Data characteristics of the AMP-AD human RNA-seq datasets.

**Table 2 -** Enrichment for heat shock response and unfolded protein response pathways for non-cell type specific modules. Fisher’s Exact Test odds ratio and adjusted p-values of gene set enrichment are shown.

## METHODS

### Study design and data collection

Details of sample collection, postmortem sample descriptions, tissue and RNA preparation, library preparation and sequencing, and sample QC are provided in previously published work (Allen et al., 2016; Jager et al., 2018; Wang et al., 2018).

### AD definition and cross-study harmonization

Sub-samples were selected to harmonize the LOAD case - control definition across the three studies for all differential expression analyses. To compare analysis results across studies and to get an understanding of LOAD biology across different tissues, we harmonized the LOAD definition across three studies. The motivation was to define LOAD cases as those with both clinical and neuropathological evidence for definitive late onset Alzheimer’s disease - i.e. a high burden of neurofibrillary tangles, neuritic amyloid plaques, and cognitive impairment with little evidence of other pathology (Jack et al., 2018). Controls were concordantly defined as patients with a low burden of plaques and tangles, as well as no evidence of cognitive impairment if available. As such, for the ROSMAP study, we had individuals with a Braak neurofibrillary tangle (NFT) score (Braak et al., 2006) greater than or equal to 4, CERAD score less than or equal to 2, and a cognitive diagnosis of probable AD with no other causes as LOAD cases, Braak less than or equal to 3, CERAD score greater than or equal to 3, and cognitive diagnosis of ‘no cognitive impairment’ as LOAD controls. For MSBB, we analogously defined LOAD cases as those with CDR score greater than or equal to 1, Braak score greater than or equal to 4, and CERAD neuritic and cortical plaque score greater than or equal to 2 as LOAD cases, and CDR scores less than or equal to 0.5, Braak less than or equal to 3, and CERAD less than or equal to 1 as LOAD controls. It is to note here that the definitions of CERAD differs between ROSMAP and MSBB studies. For the Mayo Clinic RNASeq study, cases were defined based on neuropathology, with LOAD cases being based on Braak score greater than or equal to 4 and CERAD neuritic and cortical plaque score greater than 1 whereas LOAD controls being those defined as Braak less than or equal to 3, and CERAD less than 2. Further details concerning the diagnosis in the Mayo RNASeq study have been previously published (Allen et al., 2018a).

### RNA-Seq Reprocessing, library normalization and covariates adjustment

To avoid some of the technical variabilities arising due to RNA-seq alignment and quantification, and also to account for some of the technical variabilities we reprocessed and realigned all the RNA-Seq reads from the source studies (Allen et al., 2016; Jager et al., 2018; Wang et al., 2018). The reprocessing was done using a consensus set of tools with only library type-specific parameters varying between pipelines. Picard (https://broadinstitute.github.io/picard/) was used to generate FASTQs from source BAMs. Generated FASTQ reads were aligned to the GENCODE24 (GRCh38) reference genome using STAR and gene counts were computed for each sample. To evaluate the quality of individual samples and to identify potentially important covariates for expression modeling, we computed two sets of metrics using the CollectAlignmentSummaryMetrics and CollectRnaSeqMetrics functions in Picard.

To account for differences between samples, studies, experimental batch effects and unwanted RNA-Seq specific technical variations we performed library normalization and covariate adjustments for each study separately using fixed/mixed effects modeling. The workflow consist of following steps: (i) gene filtereing: Genes that are expressed more than 1 CPM (read Counts Per Million total reads) in at least 50% of samples in each tissue and diagnosis category was used for further analysis, (ii) conditional quantile normalisation, was applied to account for variations in gene length and GC content, (iii) sample outlier detection using principal component analysis and clustering, (iv) Covariates identification and adjustment, where confidence of sampling abundance were estimated using a weighted linear model using voom-limma package in bioconductor (Ritchie et al., 2015). For most analyses, we perform a variant of fixed/mixed effect linear regression as shown here: gene expression ~ Diagnosis + Sex + covariates + (1| Donor) or gene expression ~ Diagnosis x Sex + covariates + (1|Donor), where each gene in linearly regressed independently with Diagnosis, variable explaining the AD status of an individual, identified covariates and donor information as random effect. Observation weights (if any) were calculated using the voom-limma (Ritchie et al., 2015) pipeline. So that observations with higher presumed precision will be up-weighted in the linear model fitting process. All these workflows were applied separately for each of the three studies.

### Meta-Differential Expression Analysis

All the differential and meta-differential expression analysis were performed as weighted fixed/mixed effect linear models using the voom-limma (Ritchie et al., 2015) package in R. For each gene, linear regression was fit with biological and technical covariates that were associated with the top principal components of the expression data, as identified above. Two of the three studies - MSBB and Mayo RNAseq - obtained more than one tissue from the same donors. Therefore, except ROSMAP study, donor-specific effects were explicitly modeled as random effects. Different models were built for understanding the effects of diagnosis and sex-specific diagnosis effects. Depending on the model, coefficients related to either diagnosis or diagnosis time sex was statistically tested for being non-zero, implying an estimated effect for the primary variable of interest is above and beyond any other effect from the covariates. This test produces t-statistic (then moderated in a Bayesian fashion) and corresponding p-value. P-values were then adjusted for multiple hypothesis testing using false discovery rate (FDR) estimation, and the differentially expressed genes were determined as those with an estimated FDR below, or at, 5% with a corresponding absolute expression and fold-change cutoffs. To identify genes with evidence for change in expression across studies, we next performed a meta-analysis using a random effect and fixed effect models using rmeta r package (https://cran.r-project.org/web/packages/rmeta/index.html). The random effect model was selected as a conservative approach to correct for variation across studies.

### Network Inference and Module Identification

We apply five distinct network module identification methodologies to each of the seven tissue specific expression data sets. This includes MEGENa (Song and Zhang, 2015), WINA (Wang et al., 2016), metanetwork, rWGCNA (Parikshak et al., 2016), and speakEasy (Gaiteri et al., 2015) to characterize a comprehensive landscape of transcriptomic variation across the seven brain regions and three studies. Briefly, MEGENa (Song and Zhang, 2015) is a method that infers a sparse graph based on a distance to define multiscale module definitions from coexpression data. Speakeasy is a label propagation method to identify robust coexpression modules that are identified both top up and bottom down (Gaiteri et al., 2015), rWGCNA is a version of WGCNA (Langfelder and Horvath, 2008b) that includes bootstrapping to identify robust modules, WINA is also a variation on WGCNA that includes a modified tree cutting method to identify modules (Wang et al., 2016). The metanetwork inference methodology is inspired by the DREAM5 method (Marbach et al., 2012), where ensemble inference methodologies were identified as more robust for identification of gene-gene interactions from coexpression data (Marbach et al., 2012).

### Metanetwork coexpression graph learning algorithm

We construct a statistical network of gene co-expression using an ensemble network inference algorithm. Briefly, we apply nine distinct gene co-expression network inference methodologies ARACNe (Margolin et al., 2006), Genie3 (Huynh-Thu et al., 2010), Tigress (Haury et al., 2012), Sparrow (Logsdon et al., 2015), Lasso (Krämer et al., 2009), Ridge (Krämer et al., 2009), mrnet (Meyer et al., 2007), c3net (Altay and Emmert-Streib, 2010) and WGCNA (Langfelder and Horvath, 2008b) and rank the edge lists from each method based on the method specific edge weights, identify a mean rank for each edge across methods, then identify the total number of edges supported by the data with Bayesian Information Criterion for local neighborhood selection with linear regression. The ensemble approach is inspired by work DREAM consortia (Marbach et al., 2012) showing that ensemble methods are better at generating robust gene expression networks across heterogeneous data-sets.

### Metanetwork module identification methodology

We identify metanetwork modules in each tissue type based on the inferred network topology with a consensus clustering algorithm (Wilkerson and Hayes, 2010) applied to multiple individual module identification methods. We ran individual network clustering methods applied to each of the seven network topologies. These methods included CFinder (Adamcsek et al., 2006), GANXiS (Gaiteri et al., 2015), a fast greedy algorithm (Clauset et al., 2004), InfoMap (Rosvall and Bergstrom, 2008), LinkCommunities (Ahn et al., 2010), Louvain (Blondel et al., 2008), Spinglass (Traag and Bruggeman, 2009), and Walktrap (Pons and Latapy, 2005), methods. All implementations are from the igraph package (Csardi et al.) in R.

### Aggregate module identification

For all 2978 modules identified across tissues (**Supplementary Table S1**, 10.7303/syn10309369.1), we first identify which modules are enriched for >=1 AD specific differential expressed gene set from the DEG meta-analysis (10.7303/syn11914606). This restricts the total number of individual modules to 660 that show evidence of differential expression as a function of disease status. Next, we construct a within tissue module graph using a Fisher’s exact test for pairwise overlap of gene sets between each pair of these 660 individual modules. An example of this graph is shown in **Figure S1**. We then apply the edge betweenness graph clustering method (Pons and Latapy, 2005) to identify aggregate modules from these module graphs that represent meta modules that are both differentially expressed and identified by multiple independent module identification algorithms. With this approach we identify 30 aggregate module definitions (10.7303/syn11932957.1) across the seven tissue types and three studies.

### Enrichment analyses

Aggregate modules were interpreted using functional and cell type enrichment analysis. We performed a battery of enrichment tests to understand biological functionality, including evaluating primary hypotheses previously implicated by genetic findings in AD research, performing exploratory analyses of a large number of gene sets (such as those obtained from Gene Ontology), and performing enrichment for brain tissue specific cell types. We started by curating three categories of gene sets for analyzing the differential expression data and network modules: 1) a small group of pathways and gene sets previously implicated in genome-wide genetic studies of AD (“hypothesis-driven”), 2) a collection of thousands of “hypothesis-free” gene sets from large databases like GO, Wikipathways and Reactome, that would allow us to potentially characterize novel biology arising in brain expression related to AD, and 3) Brain specific cell type markers to potentially understand the changes in various cell type fractions (Zhang et al., 2014). AMP-AD specific gene sets were constructed by taking the union of gene set definitions reported in each of the following reports: RNA-binding protein modules (Johnson et al., 2018), oligodendroglial modules from MSSM (McKenzie et al., 2017), AD vs Control oligodendroglial modules in the Mayo RNAseq study (Allen et al., 2018a), and Module 109 from the ROSMAP study (Mostafavi et al., 2018).

Genes not measured in our data are filtered from the annotated gene sets. Annotated gene sets with less than 10% of genes expressed in our data sets were removed. Fisher’s exact test was used to test enrichment of each gene set with the annotated set. Resulting p-values were corrected independently for each set using Benjamini-Hochberg method for significance testing, owing to the differences in their hypothesis. Gene sets that had a minimum overlap of at least 3 genes were considered for further interpretation.

### Statistics, code and data availability

All computation and calculations were carried out in the R language for statistical computing (version 3.3.0 - 3.5.1). Significance levels for p-values were set at 0.05 (unless otherwise specified), and analyses were two-tailed. An R package with all code for the metanetwork algorithm is available at https://github.com/Sage-Bionetworks/metanetwork, and a toolkit for integrating metanetwork with AWS high performance compute cluster cfncluster, and Synapse is available here https://github.com/Sage-Bionetworks/metanetworkSynapse. Furthermore, all code used to generate aggregate modules and figures are available in this R package: https://github.com/Sage-Bionetworks/AMPAD, with the following notebook collating the primary results: https://github.com/Sage-Bionetworks/AMPAD/blob/master/manuscript_analyses.Rmd.

## SUPPLEMENTAL INFORMATION LEGENDS

**Figure S1.**
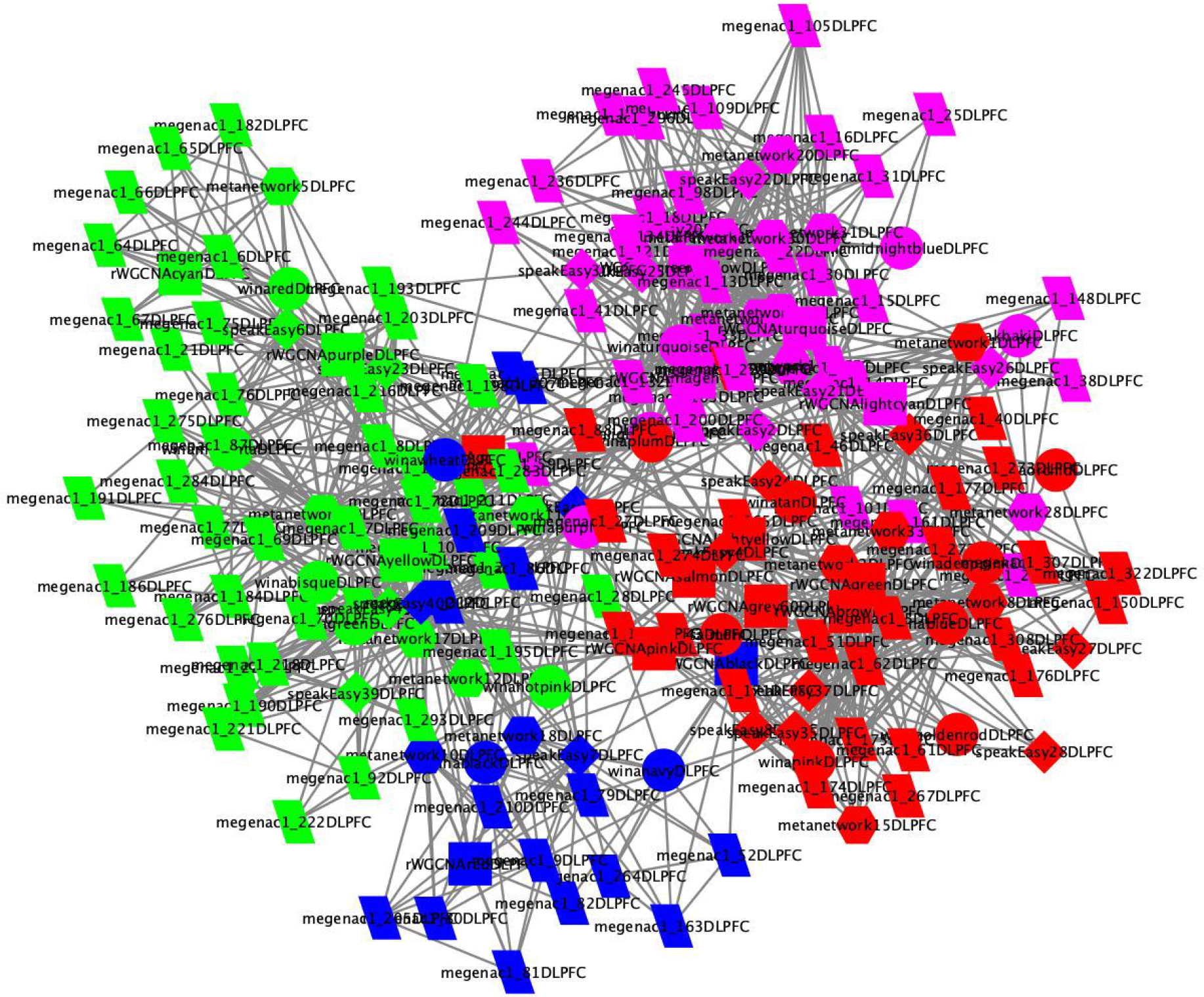
Clustering of individual AD coexpression modules. Similar coexpression structure was observed within each data set across methods, as indicated by significant overlap in module memberships. This module graph shows individual modules that are significantly enriched for at least one DEG meta-analysis signature in DLPFC (ROSMAP). Each node is a module, and an edge is drawn between modules if there is a statistically significant overlap of genes between the two modules. The edge betweenness clustering algorithm identifies four robust meta modules, which are colored green, purple, red and blue respectively.

**Table S1** - Counts of number of individual coexpression modules identified by method and brain region (10.7303/syn10309369.1).

**Table S2 –** Study demographics for each of the AMP-AD studies for samples with available bulk homogenate RNA-seq data.

**Table S3** – Module assignment to consensus clusters and module size.

**Table S4 –** Gene set enrichment results for aggregate modules compared to AMP-AD derived gene sets.

**Table S5 –** Gene set enrichment results for aggregate modules and AD gene sets against genes up-regulated in AD from the differential expressed gene sets from the random effect meta-analysis of differential expression.

**Table S6 -** Gene set enrichment results for aggregate modules and AD gene sets against genes down-regulated in AD from the differential expressed gene sets from the random effect meta-analysis of differential expression.

**Table S7 -** Gene set enrichment results for aggregate modules and AD gene sets against genes up-regulated in male AD from the differential expressed gene sets from the random effect meta-analysis of differential expression.

**Table S8 -** Gene set enrichment results for aggregate modules and AD gene sets against genes up-regulated in female AD from the differential expressed gene sets from the random effect meta-analysis of differential expression.

**Table S9 -** Gene set enrichment results for aggregate modules and AD gene sets against genes down-regulated in female AD from the differential expressed gene sets from the random effect meta-analysis of differential expression.

**Table S10 -** Gene set enrichment results for aggregate modules and AD gene sets against genes down-regulated in male AD from the differential expressed gene sets from the random effect meta-analysis of differential expression.

