## Supplementary material for "Meta-analysis of the human brain transcriptome identifies heterogeneity across human AD coexpression modules robust to sample collection and methodological approach": Tables

**Table 1**

| **Study** | **No of individuals** | **Tissue** | **RNA Library Prep** | **Strand** | **Sequencer** | **No of Samples** | **Median Read Depth (bn bases)** |
| --- | --- | --- | --- | --- | --- | --- | --- |
| MAYO | 302 | TCX | Poly-A | Double | Ilumina HiSeq 2000 | 264 | 12.58 |
|  |  | CBE |  |  |  | 263 | 13.5 |
| MSSM | 300 | FP | Ribozero | Single | Ilumina HiSeq 2000 | 260 | 3.85 |
|  |  | STG |  |  |  | 240 | 3.23 |
|  |  | PHG |  |  |  | 225 | 3.56 |
|  |  | IFG |  |  |  | 230 | 3.49 |
| ROSMAP | 632 | DLPFC | Poly-A | Double | Ilumina HiSeq | 632 | 8.36 |
| **Total** | 1234 |  |  |  |  | 2114 |  |

**Table 2**

| Consensus Cluster E Module | Attenuation phase of transcriptional response to heat shock(Abravaya, Phillips and Morimoto, 1991; Fabregat *et al.*, 2016) | Detection of unfolded protein (GO:0002235)(‘Gene Ontology Consortium: going forward’, 2015) | Response to unfolded protein (GO:0006986)(‘Gene Ontology Consortium: going forward’, 2015) | HSF1 activation (Zuo, Rungger and Voellmy, 1995; Cotto, Kline and Morimoto, 1996; Fabregat *et al.*, 2016) |
| --- | --- | --- | --- | --- |
| CBEblue | 13 (1.2x10^-5^) | 8.4 (1.5x10^-5^) | 8.4 (1.5x10^-5^) | 5.9 (1.1x10^-3^) |
| DLPFCturquoise | 8.5 (1.6x10^-4^) | 9.1 (3.9x10^-6^) | 9.1 (3.9x10^-6^) | 5.4 (3.8x10^-3^) |
| TCXbrown | 12 (7.6x10^-6^) | 5.3 (4.4x10^-3^) | 5.3 (4.4x10^-3^) | 10.5 (6.1x10^-6^) |
| STGturquoise | 11 (1.6x10^-5^) | 8.1 (1.7x10^-5^) | 8.1 (1.7x10^-5^) | 4.8 (1.1x10^-2^) |
| PHGblue | 9.5 (5.4x10^-5^) | 3.6 (3x10^-2^) | 3.6 (3x10^-2^) | 7.6 (9.8x10^-5^) |

**Table S1**

| Brain Region (Study) | Megena | metanetwork | rWGCNA | SpeakEasy | WINA |
| --- | --- | --- | --- | --- | --- |
| PHG (MSBB) | 299 | 16 | 13 | 24 | 18 |
| STG (MSBB) | 277 | 16 | 9 | 22 | 12 |
| IFG (MSBB) | 296 | 17 | 12 | 41 | 63 |
| FP (MSBB) | 274 | 17 | 16 | 35 | 51 |
| TCX (Mayo) | 352 | 33 | 24 | 49 | 44 |
| CBE (Mayo) | 340 | 33 | 19 | 59 | 47 |
| DLPFC (ROSMAP) | 291 | 34 | 19 | 58 | 48 |
| Total | 2129 | 166 | 112 | 288 | 283 |

**Table S2**

| Study (N donors) | Tissue Type | N - samples | N – females / males | N – AD / Control | AOD – mean (sd) | PMI – mean (sd) | RIN – mean (sd) | N - AD females / males | N - control females / males |
| --- | --- | --- | --- | --- | --- | --- | --- | --- | --- |
| ROSMAP (632) | DLPFC | 632 | 404 / 228 | 155 / 86 | 89 (6.6) | 7.3 (4.8) | 7.1 (1.0) | 109 / 46 | 47 / 39 |
| MayoRNAseq (302) | TCX | 264 | 136 / 128 | 80 / 73 | 80 (8.3) | 7.0 (5.2) | 8.2 (0.9) | 49 / 31 | 37 / 36 |
|  | CBE | 263 | 133 / 130 | 79 / 74 | 80 (8.3) | 7.3 (5.9) | 8.1 (0.97) | 47 / 32 | 37 / 37 |
| MSBB (300) | FP | 260 | 167 / 93 | 118 / 49 | 83 (7.4) | 7.3 (5.5) | 6.4 (1.4) | 82 / 36 | 25 / 24 |
|  | IFG | 230 | 151 / 79 | 101 / 42 | 84 (7.4) | 7.3 (5.5) | 7.5 (2.3) | 71 / 30 | 19 / 23 |
|  | PHG | 225 | 143 / 82 | 98 / 46 | 83 (7.8) | 7.2 (5.6) | 5.9 (1.8) | 68 / 30 | 23 / 23 |
|  | STG | 240 | 151 / 89 | 111 / 41 | 83 (7.6) | 6.9 (5.1) | 6.0 (1.4) | 72 / 21 | 39 / 20 |
